# Dopaminergic Ric GTPase activity impacts amphetamine sensitivity and sleep quality in a dopamine transporter-dependent manner in *Drosophila melanogaster*

**DOI:** 10.1101/2021.02.18.431857

**Authors:** Rita R. Fagan, Patrick J. Kearney, Dino Luethi, Nicholas C. Bolden, Harald H. Sitte, Patrick Emery, Haley E. Melikian

## Abstract

Dopamine (DA) is required for movement, sleep, and reward, and DA signaling is tightly controlled by the presynaptic DA transporter (DAT). Therapeutic and addictive psychostimulants, including methylphenidate (Ritalin; MPH), cocaine, and amphetamine (AMPH), markedly elevate extracellular DA via their actions as competitive DAT inhibitors (MPH, cocaine) and substrates (AMPH). DAT silencing in mice and invertebrates results in hyperactivity, reduced sleep, and blunted psychostimulant responses, highlighting DAT’s essential role in DA-dependent behaviors. DAT surface expression is not static; rather it is dynamically regulated by endocytic trafficking. PKC-stimulated DAT endocytosis requires the neuronal GTPase, Rit2, and Rit2 silencing in mouse DA neurons impacts psychostimulant sensitivity. However, it is unknown whether or not Rit2-mediated changes in psychostimulant sensitivity are DAT-dependent. Here, we leveraged *Drosophila melanogaster* to test whether the *Drosophila* Rit2 ortholog, Ric, impacts dDAT function, trafficking, and DA-dependent behaviors. Orthologous to hDAT and Rit2, dDAT and Ric directly interact, and the constitutively active Ric mutant Q117L increased dDAT surface levels and function in cell lines and *ex vivo Drosophila* brains. Moreover, DAergic RicQ117L expression caused sleep fragmentation in a DAT-dependent manner but had no effect on total sleep and daily locomotor activity. Importantly, we found that Rit2 is required for AMPH-stimulated DAT internalization in mouse striatum, and that DAergic RicQ117L expression significantly increased *Drosophila* AMPH sensitivity in a DAT-dependent manner, suggesting a conserved impact of Ric-dependent DAT trafficking on AMPH sensitivity. These studies support that the DAT/Rit2 interaction impacts both baseline behaviors and AMPH sensitivity, potentially by regulating DAT trafficking.

## Introduction

Dopamine (DA) is critical for a variety of behaviors, including motor function, sleep, learning, and reward^1,2^. Multiple neuropsychiatric and neurologic disorders are linked to DAergic dysfunction, including addiction, attention-deficit hyperactivity disorder (ADHD), autism spectrum disorder (ASD), schizophrenia, and Parkinson’s disease (PD)^3-7^. Extracellular DA levels are tightly restricted by the presynaptic DA transporter (DAT). Following evoked DA release, DAT takes up released DA back into DAergic presynaptic terminals, thereby limiting the spatial and temporal DA signal^8^. DAT not only limits extracellular DA levels, but is also critical to maintain homeostatic DA levels and synaptic DAergic tone across multiple species^9^. DAT’s pivotal role in regulating DAergic signaling is best illustrated by observing the consequences that follow from either pharmacological DAT inhibition or genetic DAT perturbation. For example, addictive and therapeutic psychostimulants, such as cocaine, AMPH, methamphetamine (METH), and methylphenidate (Ritalin, MPH), are competitive DAT antagonists (cocaine, MPH) and substrates (AMPH, METH) that increase extracellular DA levels and drive hyperlocomotion, sleep perturbation, and rewarding behaviors in both vertebrates and invertebrates^10,11^. Moreover, genetic DAT ablation in both vertebrates and invertebrates 1) blocks psychostimulant-induced hyperactivity and reward, and 2) causes hyperactivity and sleep loss^12-15^. DAT knock-out (*DAT*^*-/-*^) mice display alterations in their sleep and waking patterns in which their waking episodes (or bouts) are three times longer than wildtype controls^13^. *Drosophila melanogaster* DAT (dDAT)-null fruit flies (“fumin”, *fmn*) also sleep significantly less than wildtype flies, with shorter and fewer resting bouts^14^. In addition to mouse and *Drosophila* DAT manipulations, numerous DAT coding variants have been reported in ADHD, ASD, and PD patients^16-21^, underscoring that DAT availability and function play an indispensable role in basal DAergic function.

Given DAT’s fundamental role in DAergic signaling, the cellular mechanisms that modify DAT function and/or expression at the plasma membrane are ideally poised to impact DAergic neurotransmission and DA-dependent behaviors. Decades of research support that DAT surface expression is acutely modulated by endocytic trafficking^22-24^. Both PKC activation and AMPH exposure acutely increase DAT internalization, resulting in a rapid net loss of DAT from the plasma membrane, as demonstrated both *in vitro* and in *ex vivo* mouse striatal slices^8,22,24-27^. However, it remains unknown whether or not regulated DAT trafficking impacts DA-dependent behaviors.

Rit2 is a neuronal-specific^28-30^ GTPase that is enriched in DA neurons^31^, and is highly homologous to Rit1, which is ubiquitously expressed^32,33^. Multiple genome-wide association studies (GWAS) have identified Rit2 as a risk allele selectively in DA-related neuropsychiatric disorders, including PD, ASD, schizophrenia, and speech delay^34-42^. We previously reported that Rit2 binds directly to DAT^43,44^, and that Rit2 is required for PKC-stimulated DAT endocytosis in both cell lines and striatal DAergic terminals^43,44^. PKC activation drives DAT and Rit2 to dissociate at the cell surface, and failure of DAT and Rit2 to dissociate correlates with increased DAT stability at the plasma membrane, and inability to undergo PKC-stimulated endocytosis^43^. Importantly, we also found that conditional Rit2 knockdown (Rit2-KD) in DA neurons significantly increases acute cocaine sensitivity *in vivo* in male mice^45^. These data raise the possibility that Rit2-mediated changes in psychostimulant sensitivity are due to inability of DAT to undergo regulated trafficking. However, it is not known whether Rit2-dependent behavior phenotypes in mice are DAT-dependent or DAT-independent. In order to address this question, we leveraged the model organism, *Drosophila melanogaster*, which expresses a single Rit2 homolog, Ric. Ric shares ∼71% homology with Rit2, but is expressed ubiquitously in the fruit fly, similar to Rit1. Given that the DA system is highly conserved between flies and mammals, *Drosophila* provide a highly tractable and simplified model to study the impact of regulated DAT trafficking on DA-dependent behaviors^46^. Similar to *DAT*^*(-/-)*^ mice, DAT null flies (*fmn)* are hyperactive, lack psychostimulant response, and numerous reports indicate that hDAT suffices to rescue hyperactivity and psychostimulant response in *fmn* flies^15,21,47,48^. Moreover, expressing hDAT ASD and ADHD coding variants on the *Drosophila fmn* background leads to hyperactivity and disrupted psychostimulant response^21,49^. In the present study, we found that Ric GTPase binds to and regulates dDAT function and expression, similar to its mammalian homolog, Rit2. Moreover, DAergic expression of the constitutively active RicQ117L leads to reduced sleep consolidation and increased AMPH sensitivity in a DAT-dependent manner.

## MATERIALS AND METHODS

### Materials

Nisoxetine was from Tocris-Cookson. Rabbit anti-HA antibody (clone C29F4) was from Cell Signaling Technology and rat anti-DAT antibody (MAB369) was from Millipore. HRP-conjugated goat anti-rabbit (Cat#111-035-003) and goat anti-rat (Cat#112-035-062) secondary antibodies were from Jackson ImmunoResearch. All other reagents were from either Sigma-Aldrich or Fisher Scientific and were of the highest possible grade.

### cDNA plasmids

dDAT and WT-Ric plasmids were from Addgene and Dr. Douglas Harrison (University of Kentucky), respectively, were subcloned into pcDNA3.1(+), and HA tags were inserted at their N-termini by PCR. Ric S73N and Q117L mutants were generated using the QuikChange II mutagenesis kit (Stratagene). YFP- and CFP-dDAT and Ric constructs were generated by subcloning into pEYFP-C1 and pECFP-C1 vectors, respectively, at the HindIII and XbaI sites. All sequences were verified by Sanger sequencing (GeneWiz).

### Fly stocks

All fly stocks were maintained on low yeast medium as previously described^50,51^. Transgenic strains *TH-GAL4* and *fmn* were gifts from Dr. Heinrich Matthies (University of Alabama School of Medicine). *UAS-HA-Ric, UAS-HA-RicQ117L*, and *UAS-HA-RicS73N* lines were generated by subcloning into the 5xUAS, w+ vector^52,53^ (gift of Dr. Marc Freeman, Vollum Institute), and were inserted into the genome via phiC31-mediated integration (BestGene). All fly strains were backcrossed to the isogenic *w*^*1118*^ wildtype (+/+; VDRC60000) strain for at least six generations and were balanced prior to behavioral studies. Transgenes were verified by eye color or PCR.

### Drosophila circadian behavior and feeding

Sleep and locomotor behavior were assessed in male progeny 1-3dpe using the *Drosophila* Activity Monitor system, as previously described^50,51,54^. 1-3 days old male flies were loaded in Trikinetics DAM monitors (Waltham, MA) and exposed to 6 days of LD cycles. The last three days were used to measure sleep. At least 3 independent experiments, with multiple flies/experiment, were performed, and data from 3 days of LD were averaged for each experiment. Data were analyzed using the sleep and circadian analysis MATLAB program (SCAMP)^55^. A sleep bout was defined as 5 consecutive minutes of inactivity. For AMPH feeding studies, flies were starved and then fed either base fly behavior food, or food supplemented with the indicated AMPH doses, as previously reported^48,49^. Locomotor activity was recorded for 24 hrs after introducing AMPH- or vehicle-supplemented food. For caffeine and carbamazepine (CBZ) studies, flies were not starved, and locomotor activity was recorded for either 24 hrs immediately (caffeine) or 24 hours (CBZ) following food change to either vehicle- or drug-supplemented sucrose-agar food. For CBZ studies, 20 mg/ml CBZ stocks were prepared in 45% (2-hydroxypropyl)-β-cyclodextrin (Sigma) and mixed into the standard agar medium. (2-hydroxypropyl)-β-cyclodextrin vehicle controls were prepared in parallel. Flies that died during the assay were excluded from further analysis.

### Cell culture and transfections

HEK293T and HEK293 (FRET studies) cells were maintained at 37°C, 5% CO_2_, and transiently transfected with Lipofectamine or JetPrime (FRET studies), as previously described^43^. Cells were seeded one day prior to transfection onto poly-D-lysine coated plates at a density of 5×10^5^ cells/well for biochemical studies (6-well), or 1.67×10^4^ cells/well for [^3^H]DA uptake studies (96-well white opaque/clear bottom plates), and were transfected using 2μg plasmid DNA/well for 6 well dishes, and 0.067μg plasmid DNA/well for 96 well plates, at a lipid:DNA ratio of 2:1. For co-transfection studies, cDNA was transfected at a co-expressor:DAT ratio of 4:1. For FRET microscopy, 2×10^4^ cells/chamber were seeded in an ibidi 8-well chambered coverslips and were transfected using 0.25μg plasmid DNA/chamber. All assays were conducted 24-48 hrs post-transfection.

### Live cell FRET microscopy

FRET was measured and at the plasma membrane and analyzed as previously described^43^. Normalized FRET (NFRET) was calculated as follows:

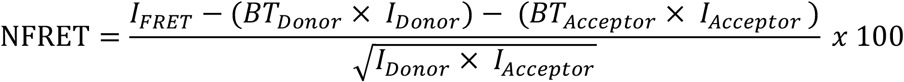

A fused CFP-YFP construct (CYFP) was included as an internal instrument control in each experiment, and non-fused donor and acceptor fluorophores were included as a negative control. N values indicate the number of independent cells imaged per condition, over 2 independent experiments.

### [^3^H]DA and [^3^H]5-HT uptake

#### Transfected cells

[^3^H]DA uptake was measured in cells transfected with either dDAT, hSERT, or dSERT essentially as previously described^27,56,57^. Triple technical replicates were assessed and averaged for each independent experiment. For methylphenidate dose-response curves, cells were incubated with the indicated methylphenidate concentration for 30 min, 37°C, prior to initiated transport with either 50nM [^3^H]DA or [^3^H]5-HT. For AMPH dose-response curves, 50nM [^3^H]DA was spiked with the indicated AMPH concentrations, such that both AMPH and [^3^H]DA were added simultaneously to initiate uptake. Non-specific uptake was defined as follows for the indicated transporters: dDAT: 10µM nisoxetine (a highly potent and selective dDAT inhibitor^58^); hDAT: 10µM GBR12909; hSERT: and dSERT:. HA-Ric expression was confirmed in each experiment via immunoblotting parallel transfected cells.

#### Ex vivo Drosophila brains

Whole brain uptake assays were performed as described previously, with slight modifications^49^. Brains from male progeny 0-3dpe were dissected in hemolymph-like solution (HL3; 70mM NaCl, 5mM KCl, 1.5mM Ca^2+^ acetate, 10mM MgSO_4_, 5mM HEPES, supplemented daily with fresh 115mM sucrose, 5mM trehalose, 10mM NaHCO_3_, 10µM sodium ascorbate, 10µM pargyline), and 4 brains/condition were pooled and used for a single measurement in each independent experiment. Brains were pre-incubated in HL3 ±10µM nisoxetine for 20min at 25°C in cell culture inserts, and DA uptake was initiated by adding 1µM [^3^H]DA. Uptake proceeded for 7 min, RT, and was terminated by rapidly washing brains 3×1mL ice cold HL3. Brains were transferred to 96-well white/clear plates, solubilized in scintillation fluid, and radioactivity was measured using a Wallac MicroBeta scintillation plate counter.

### Cell surface biotinylation

Surface dDAT was measured by biotinylation in transfected HEK293T cells as previously described for hDAT^27,43,57^. Biotinylated proteins were quantitatively recovered by streptavidin batch chromatography from equivalent amounts of cellular protein, and were eluted in 2X Laemmli sample buffer, 30 min, room temperature with rotation. Eluted proteins and input lysates were resolved by SDS-PAGE and proteins were detected and quantified by immunoblotting. Immunoreactive bands were detected by chemiluminescence (SuperSignal™ West Dura Extended Duration Substrate, ThermoFisher) and were captured with a CCD-camera (VersaDoc, Biorad). Bands in the linear range of detection were quantified using Quantity One software (Bio-rad), and the %surface DAT was determined by normalizing surface DAT to its corresponding DAT input signal. Single values were obtained in each independent experiment.

### Vertebrate Animals

All studies were conducted in accordance with UMASS Medical School IACUC Protocol 202100046 (formerly A-1506) (H.E.M). *Pitx3*^*IRES2-tTA*^ mice on the *C57Bl/6* background were the generous gift of Dr. Huaibin Cai (National Institute on Aging) and were continually backcrossed to *C57Bl/6J* mice. Mice were maintained in 12hr light/dark cycle at constant temperature and humidity and food and water were available ad libitum. Both male and female mice were used equivalently.

### AAV production and stereotaxic viral delivery

AAV9 particles expressing TRE-eGFP and TRE-shRit2-eGFP were produced by the UMASS Medical School Viral Vector Core, as previously described^43,45^. P21-25 *Pitx3*^*IRES2-tTA*^ mou VTA were bilaterally injected with 1µl AAV particles (Bregma: anterior/posterior: -3.08mm, medial/lateral: ±0.5mm, dorsal/ventral: -4.5mm) at a rate of 0.2µL/min as previously described^45^. Studies were performed 4-6 weeks post-injection, and viral expression was confirmed by visualizing GFP expression in midbrain slices.

### Ex vivo slice biotinylation

Mouse striatal slices were prepared and hemisected along the midline as previously described^27,43,45^, and were treated ±10µM AMPH in ACSF, 30min, 37°C with constant oxygenation, using paired contralateral hemi-slices as a vehicle and drug-treated samples. Following drug treatment, surface proteins were labeled by biotinylation, hemi-slices were subdissected to enrich for dorsal striatum, and tissue was solubilized as previously described^43,45^. Biotinylated proteins were isolated by batch streptavidin chromatography from equivalent amounts of protein, and DAT was detected by immunoblot. %Surface DAT was calculated by normalizing surface signals to their corresponding total DAT input signals, detected in parallel. Note that slice biotinylation data in the current study were acquired during the course of a previous study, in which we first reported the effect of DAergic Rit2 KD on DAT basal surface levesls^45^. To avoid duplicate reporting basal DAT surface levels from the previous study, we used those vehicle-treated DAT surface signals to determine the magnitude of AMPH’s effect on DAT surface expression in their contralateral hemislices. Therefore, data were reported as %change in DAT surface levels, normalized to vehicle-treated hemi-slices, rather than as raw surface levels. There were no statistically significant differences between male and female DAT surface levels, in either eGFP- or shRit2-injected mice. Therefore, male and female data were pooled. Data were collected from 6 independent mice per virus.

### Statistical analysis

Sample sizes were set at a minimum of n=3 for cell-based studies, n=6 for *ex vivo* biochemical studies, and n=20 for *Drosophila* behavior, based on previous studies from our laboratories indicating that these sample sizes provide sufficient power. All data were analyzed using GraphPad Prism software. Statistical outliers were identified with Grubb’s or Rout’s outlier tests (α- and Q-value set to 0.05 and 5%, respectively), and were excluded from further analysis. Student’s *t* test was used to compare between two groups, and Welch’s correction was applied in instances where standard deviations significantly differed from each other (as determined using Bartlett’s test). One- or two-way ANOVA were used to compare among more than two experimental groups, and appropriate post-hoc multiple comparisons tests were applied to determine significant differences among experimental groups, as described within each figure legend.

## RESULTS

### Ric interacts with dDAT and impacts dDAT activity

Although hDAT and Rit2 interact at the plasma membrane, it is unknown whether dDAT and Ric interact in an orthologous fashion. Therefore, we first used live-cell FRET to directly test this possibility in HEK293 cells co-transfected with CFP-dDAT and YFP-Ric. As previously reported^43,44^, CFP-hDAT/YFP-Rit2 elicited a significant FRET signal over control cells transfected with soluble CFP and YFP (Fig. 1A). CFP-dDAT and YFP-Ric likewise elicited a FRET signal that was significantly larger than CFP/YFP alone, and which did not significantly differ from the CFP-hDAT/YFP-Rit2 FRET signal (p=0.53, Fig. 1A). Given our previous results that indicate an integral role of the DAT-Rit2 interaction in regulating DAT endocytosis, these data strongly suggested that *Drosophila* Ric may similarly regulate dDAT.

**Figure 1:**
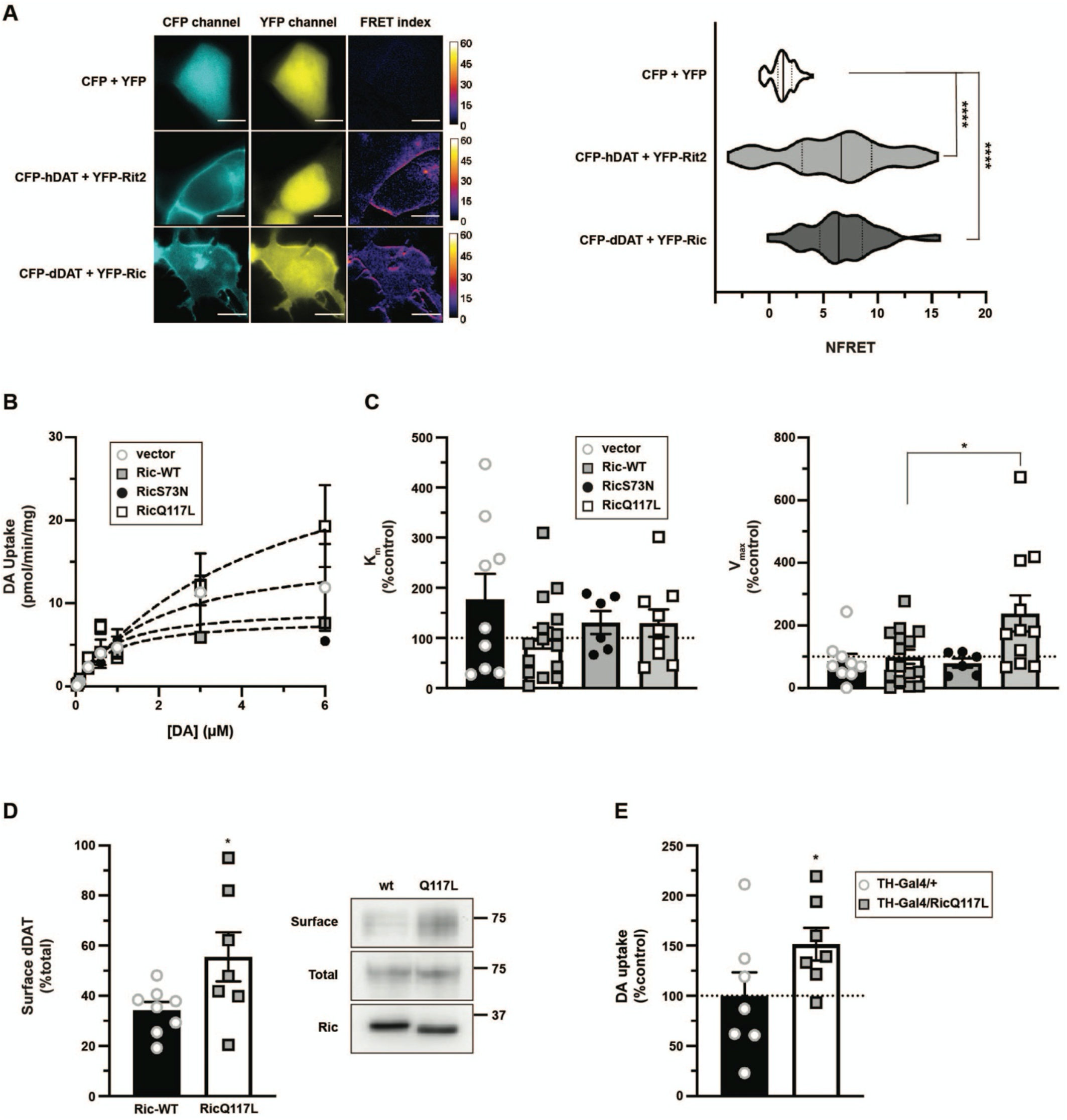
dDAT and Ric directly interact, and RicQ117L increases dDAT function and surface expression. (**A)** *Live cell FRET microscopy*. HEK293 cells were transfected with the indicated constructs and live cell FRET microscopy was performed as described in *Methods. Left:* Representative images for each indicated pair are shown in the CFP and YFP channel. The FRET index was computed after background subtraction. Scale bar = 10µm. *Right:* Average data presented as violin plots indicating the median and quartiles. Asterisks indicate a significant FRET signal as determined by Kruskal-Wallis test (p<0.0001), as compared to soluble CFP/YFP negative controls (n=77) for all FRET pairs (Dunn’s multiple comparisons test): CFP-hDAT/YFP-Rit2 (****p<0.0001, n=73) and CFP-dDAT/YFP-Ric (****p<0.0001, n=55). CFP-hDAT/YFP-Rit2 and CFP-dDAT/YFP-Ric were not significantly different from each other (p=0.53). (**B, C)**. *DA uptake kinetics*. HEK293T cells were co-transfected with HA-dDAT, and either pcDNA3.1(+) (vector), HA-Ric-WT, HA-RicQ117L, or HA-RicS73N, and [^3^H]DA uptake kinetics were measured as described in *Methods*. (**B)** Representative DA transport curves. (**C)** *Average DA uptake kinetic values*. K_m_ (*left*) and V_max_ (*right*) are expressed as a percentage of Ric-WT co-transfected controls ±S.E.M., normalized within each independent experiment. dDAT K_m_ was not significantly affected by co-transfection with either WT-Ric (n=15), RicS73N (n=6), or RicQ117L (n=9), as compared to dDAT/vector (n=9) co-transfected cells (p=0.36, one-way ANOVA, F_(3, 35)_ = 2.428). HA-RicQ117L significantly increased dDAT Vmax as compared to WT-Ric controls (one-way ANOVA: F_(3, 37)_ = 2.474, p=0.01; *p=0.01, Dunnett’s multiple comparisons test). (**D)** *Surface biotinylation*. HEK293T cells were co-transfected with HA-dDAT, and either HA-WT-Ric (n=6) or HA-RicQ117L (n=5), and dDAT surface expression was measured by surface biotinylation as described in *Methods. Right:* Representative immunoblots. *Left:* Average surface dDAT values expressed as %total dDAT ±S.E.M. (*p=0.038, one-tailed Student’s *t* test with Welch’s correction). **E**. *Ex vivo whole brain [*^*3*^*H]DA uptake*. TH-Gal4 flies were crossed to +/+ (*TH-Gal4/+*), or UAS-RicQ117L (*TH-Gal4/UAS-RicQ117L*) flies. DA uptake was measured from dissected brains as described in *Methods*. DA uptake is expressed as %TH-Gal4/+ control values ±S.E.M. (*p=0.04, one-tailed Student’s t test, n=7).

### Constitutively active Ric increases dDAT function and surface levels

We next used Ric GTPase mutants to test whether Ric activity impacts dDAT-mediated DA uptake kinetics in HEK293T cells co-transfected with dDAT and either vector, wildtype Ric (Ric-WT) constitutively active (CA) Ric (RicQ117L), or dominant negative (DN) Ric (RicS73N) (Fig. 1B). Neither WT Ric, nor any of the mutant Rics, significantly affected dDAT K_m_ values, as compared to vector co-transfected controls (Fig. 1C *top*). Similarly, neither Ric-WT nor RicS73N significantly affected dDAT V_max_ as compared to vector co-transfected cells (Fig. 1C *bottom)*. However, RicQ117L significantly increased dDAT V_max_ (Fig. 1C *bottom*). To test whether increased dDAT V_max_ was due to increased dDAT surface expression, or enhanced substrate turnover rates, we measured dDAT plasma membrane expression by surface biotinylation. dDAT surface levels were significantly increased when co-expressed with RicQ117L, as compared with cells co-transfected with Ric-WT (Fig. 1D), indicating that RicQ117L-mediated increased dDAT V_max_ was likely due to enhanced dDAT surface expression. We further tested whether RicQ117L similarly increases DA uptake *in situ*, as measured by *ex vivo Drosophila* whole brain [^3^H]DA uptake^49^. *TH-GAL4* flies were crossed to either wildtype (+/+) or *UAS-HA-RicQ117L* flies to drive RicQ117L expression selectively in DA neurons^59^, and DA uptake was measured in male progeny (*TH-GAL/UAS-RicQ117L*). RicQ117L overexpression significantly increased DA uptake as compared to control *TH-GAL4/+* flies (Fig. 1E). Together, these data demonstrate that Ric GTPase activity regulates dDAT function in intact fly brains, likely by regulating dDAT surface expression, and suggests that the dDAT/Ric interaction is orthologous to the hDAT/Rit2 interaction.

### Manipulating DAergic Ric activity does not impact locomotor activity or total sleep

Given that DA plays a central role in mammalian and *Drosophila* locomotion and sleep^60,61^, and that Ric activity impacts DAT function, we asked whether DAergic Ric activity impacts *Drosophila* basal locomotion and sleep. Using the *TH-GAL4* driver, we overexpressed Ric-WT (*TH-GAL4/UAS-Ric-WT*), DN-RicS73N (*TH-GAL4/UAS-RicS73N*), or CA-RicQ117L (*TH-GAL4/UAS-RicQ117L*) in DA neurons and monitored male fly circadian behavior. Total locomotor activity was not significantly affected in *TH-GAL4/UAS-Ric-WT, TH-GAL4/UAS-RicS73N*, or *TH-GAL4/UAS-RicQ117L* flies, compared to *TH-Gal4/+* controls, assessed over either 24 hrs, or specifically during the day or night (Fig. S1). Total sleep time was likewise unaffected, with the exception of Ric-WT overexpression which resulted in a small, but significant, decrease in daytime sleep and increase in nighttime sleep (Fig. S2).

### Increasing DAergic Ric activity decreases sleep bout consolidation in a DAT-dependent manner

Sleep and activity bout fragmentation is indicative of reduced sleep consolidation and quality, and can negatively impact health and cognitive function^62,63^. Therefore, we also analyzed our locomotor data to ask whether DAergic Ric activity impacts the number of sleep episodes. DAergic Ric-WT and DN RicS73N overexpression had no significant impact on overall (24-hr) sleep bout number, although a subtle reduction was observed during the day with WT and at night with RicS73N (Fig. 2A-B). In contrast, DAergic RicQ117L overexpression led to a significant increase in overall sleep episode number, which was entirely caused by increased daytime sleep fragmentation (Fig. 2C). This indicates a specific defect in sleep consolidation caused by DAergic RicQ117L overexpression. RicQ117L-mediated decreased sleep consolidation may be a result of the increased DA reuptake caused by overexpression, or may be due to a Ric-mediated, DAT-independent mechanism. To discern between these possibilities, we performed epistasis studies in which both driver (*TH-GAL4*) and responder (*UAS-HA-RicQ117L*) transgenes were introduced into the dDAT null background (*fmn; TH-GAL4/UAS-RicQ117L*). We predicted that if any of the locomotor phenotypes caused by RicQ117L expression were DAT-dependent, they would be lost in the DAT null background. In the absence of dDAT, RicQ117L had no effect on sleep bout number as compared to control flies (Fig. 2D), indicating that the RicQ117L-mediated sleep phenotype is DAT-dependent, and may be due to the enhanced dDAT function that we observed in intact fly brains (Fig. 1E). Alternatively, given that *fmn* flies already exhibit an increase in sleep bout frequency, there is a possibility that a ceiling effect obscured our ability to detect a DAT-independent effect of RicQ117L on sleep consolidation. To test for this possibility, we treated flies with carbamazepine (CBZ), which was reported to alter *Drosophila* sleep bout numbers in a dose-dependent manner during the 24 hours of CBZ feeding^64^. In that study, sleep episode frequency tended to increase with a low CBZ dose (0.2mg/ml), while it strongly decreased with a high dose (1.2 mg/ml). In our hands, there was no significant effect on sleep bout frequency with either 0.2mg/ml or 1mg/ml CBZ during the first 24 hours of feeding (not shown). During the second 24 period of feeding (from 24-48 hours CBZ feeding), 0.2mg/ml CBZ still had no effect on either *wt* or *fmn* sleep bout frequency. However, 1mg/ml CBZ significantly increased *fmn* sleep bout frequency and strongly trended to increase *wt* fly sleep bout frequency during the 24-48 hour period (Fig. S3). Thus, it is highly unlikely that a ceiling effect is confounding our interpretation of the epistasis studies, and RicQ117L effects on sleep consolidation are likely mediated via the effects of RicQ117L on DAT surface expression and activity.

**Figure 2.**
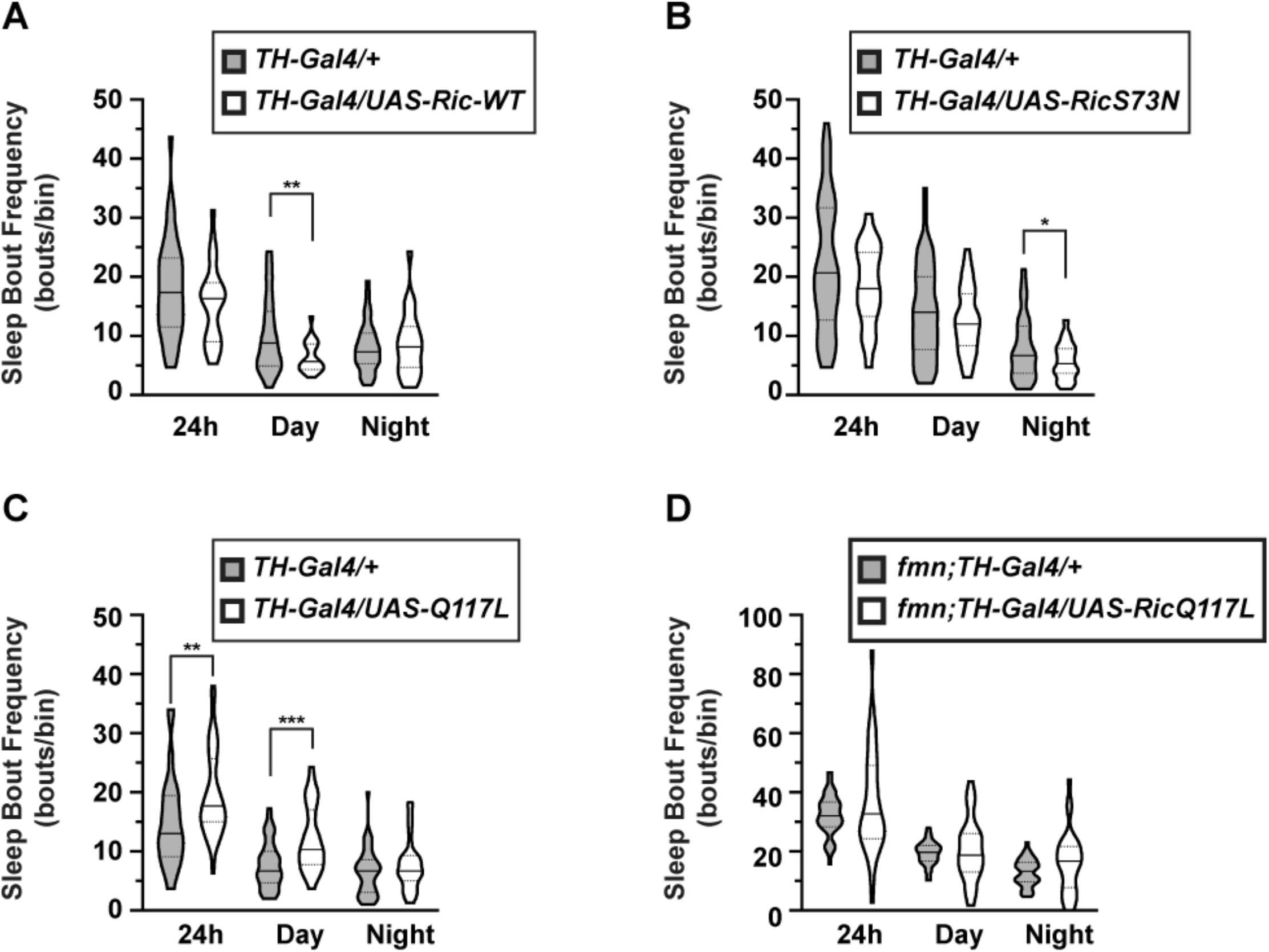
DAergic Ric activity modulates sleep bout frequency in a dDAT-dependent manner. *TH-Gal4* and *Ric-WT, UAS-RicS73N*, or *UAS-RicQ117L* flies were crossed and the number of sleep episodes in male progeny were counted for bins of 24h, daytime (12h lights-on), and nighttime (12h lights-off), and data were analyzed as described above. Data are presented as violin plot with median and quartiles indicated. (**A**) Ric-WT (n=24) had no effect on sleep bout number over 24h or at night as compared to control *TH-Gal4* flies (n=38), but significantly decreased sleep frequency during the daytime (**p=0.003, two-tailed Student’s *t* test with Welch’s correction). (**B**) RicS73N (n=41)had no effect on sleep bout number over 24h or during the day as compared to control *TH-Gal4* flies (n=47), but significantly decreased sleep frequency during at night (*p=0.02, two-tailed Student’s *t* test with Welch’s correction). (**C**) RicQ117L (n=36) significantly increased sleep bouts during the 24h bin (**p=0.005, two-tailed Student’s *t* test,) and during the day (***p=0.0001, two-tailed Student’s *t* test with Welch’s correction) as compared to control *TH-Gal4* flies (n=44), but did affect sleep bout numbers at night. (**D**) *Epistasis studies: TH-Gal4* and *UAS-RicQ117L* flies were crossed onto the dDAT-null background (*fmn*) and locomotor activity and sleep were measured in male progeny (*fmn; TH-Gal4/UAS-RicQ117L*). *fmn; TH-Gal4/UAS-RicQ117L* flies (n=31) did not exhibit a significant difference in the number of sleep episodes as compared to *fmn; TH-Gal4/+* controls (n=47) (24h: p=0.19; day: p=0.61; night: p=0.087, two-tailed Student’s *t* test with Welch’s correction).

### Ric activity impacts amphetamine sensitivity in a DAT-dependent manner

We previously reported that DAergic Rit2 knockdown significantly increases acute cocaine sensitivity in male mice^45^. However, it is not known whether Rit2-mediated changes in psychostimulant sensitivity are mediated by a DAT-dependent, or -independent, mechanism. Given that the dDAT/Ric interaction is orthologous to hDAT/Rit2, we leveraged *Drosophila* to test whether 1) Ric activity similarly impacts psychostimulant sensitivity, and 2) Ric-dependent psychostimulant sensitivity is DAT-dependent. Given the inherent challenges in administering cocaine to flies, which must be volatized, we first considered feeding flies methylphenidate (Ritalin, MPH), a monoamine transporter inhibitor with selectivity for mammalian NET and DAT over SERT. Given that flies do not express a NET, we reasoned that we might selectively inhibit DAT over SERT using MPH *in vivo*. We first tested whether MPH selectivity for mammalian DAT vs. SERT was conserved in *Drosophila* DAT and SERT by performing dose-response curves in HEK293T cells transfected with either hDAT, hSERT, dDAT, or dSERT. Results are shown in Figure 3A. While hDAT exhibited its known sensitivity to MPH (IC_50_ = 0.23 ±0.12µM, n=6), dDAT was significantly less sensitive to MPH than hDAT by more than two orders of magnitude (IC_50_ = 43 ±12.6µM, n=5), and the MPH IC_50_ measured for dDAT was comparable to that obtained for both hSERT (IC_50_ = 158 ±142µM, n=3) and dSERT (IC_50_ = 38.4 ±10.6µM, n=5). Given these results, we instead chose to assess AMPH-induced hyperlocomotion via a feeding assay^65^. AMPH elicited significant, dose-dependent, hyperlocomotion in control (*TH-GAL4/+*) flies (Fig. 3B), consistent with previous studies^65^. Surprisingly, RicQ117L expression in DAergic neurons significantly increased fly sensitivity to AMPH, resulting in significant hyperactivity at lower doses than of that required for TH-Gal4/+ controls (Fig. 3C). The increased fly AMPH sensitivity is likely due to RicQ117L effects on DAT surface stability and function. Alternatively, the RicQ117L mutant could be influencing AMPH sensitivity in a DAT-independent manner. To discern between these possibilities, we again performed an epistasis study, and expressed RicQ117L in the DAT null (*fmn)* background, to test whether the enhanced AMPH sensitivity we observed in the *TH-Gal4/UAS-RicQ117L* flies is DAT-dependent. AMPH did not significantly increase locomotion in *fmn* flies (Fig. 4A), consistent with the known requirement for DAT in AMPH-stimulated locomotion. Moreover, AMPH significantly decreased locomotion in *fmn* flies at the highest AMPH doses tested (Fig. 4A). When RicQ117L was expressed on the *fmn* background, AMPH failed to significantly increase locomotion at any of the doses tested (Fig. 4B), consistent with a DAT-dependent mechanism. Given that *fmn* flies are hyperactive at baseline, failure of AMPH to elicit an increase in locomotion in *fmn; TH-Gal4/UAS-RicQ117L* flies could potentially be due to a ceiling effect on *fmn* fly locomotion, rather than due to a DAT-dependent mechanism. Caffeine was previously reported to decrease sleep in female *fmn* flies^66^, raising the possibility that *fmn* fly locomotion may reciprocally increase in response to caffeine. To test for this possibility, we measure locomotion in response to caffeine feeding (0.5mg/ml) in male flies. As shown in Fig. 4C, caffeine significantly increased locomotion over 24 hours in both *wt* and *fmn* flies. Thus, *fmn* flies are not subject to a ceiling effect on their locomotion, and it is therefore likely that the increased AMPH sensitivity observed in the *TH-Gal4/UAS-RicQ117L* flies is mediated by the DAT-Ric interaction.

**Figure 3.**
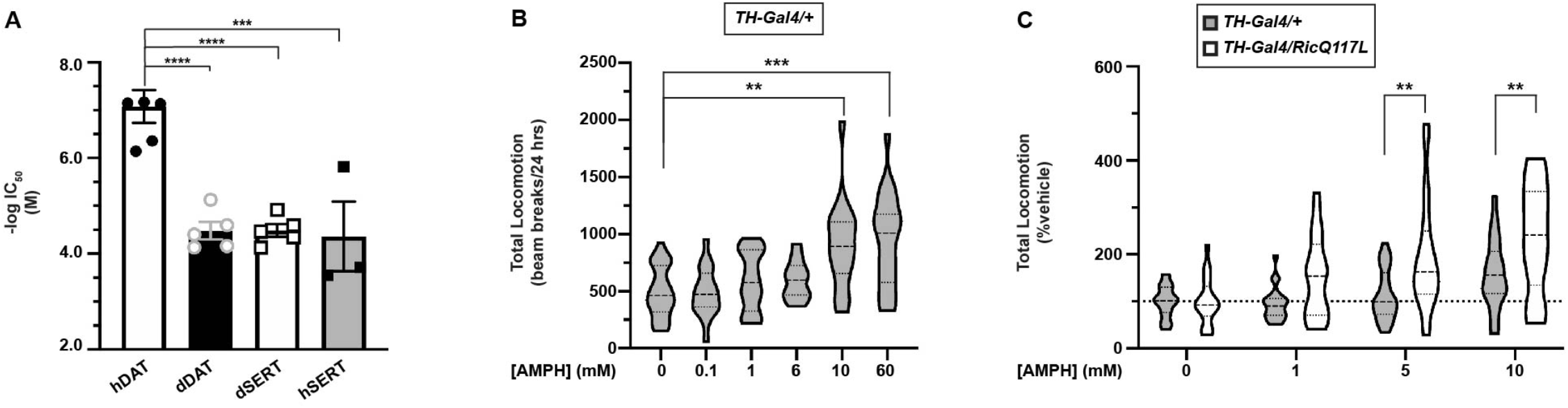
DAergic RicQ117L increases AMPH sensitivity in *Drosophila*. **(A)** *Methylphenidate potency at fly and human DAT and SERT*. HEK293T cells were transfected with cDNAs encoding either hDAT (n=7), dDAT (n=5), hSERT (n=3) or dSERT (n=5), and methylphenidate potency was measured by competition with either [^3^H]DA or [^3^H]5-HT as described in *Methods*. Methylphenidate potencies are reported as -log(IC_50_) ±S.E.M. for each transporter. Asterisks indicate a significant difference as compared to hDAT (one-way ANOVA, F_(3, 15)_ = 18.32, p<0.0001. ***p=0.0002, ****p<0.0001, Dunnett’s multiple comparisons test). **(B, C)** *Locomotor response to AMPH feeding*. (**B**) *TH-Gal4* flies were crossed to *+/+* controls, and locomotor activity was measured in progeny for 24h using food supplemented with either vehicle (n=21), or 0.1mM (n=13), 1.0mM (n=13), 10mM (n=16), or 60mM (n=13) AMPH, as described in *Methods*. Data are presented as violin plots with median and quartiles indicated. AMPH significantly increased locomotion at 10mM and 60mM doses (Kruskal-Wallis test, p<0.0001; **p=0.001, ***p=0.0002, Dunn’s multiple comparisons test, n=11-28). (**C**) *TH-Gal4/UAS-RicQ117L response to AMPH, TH-Gal4/+* (vehicle: n=26; 1.0mM: n=26; 5.0mM: n=26; 10mM: n=28) or *TH-Gal4/UAS-RicQ117L* (vehicle: n=20; 1.0mM: n=21; 5.0mM: n=24; 10mM: n=21) and AMPH-dependent locomotion was measured in progeny over 3 independent experiments. Data are presented as %vehicle-fed response, measured in parallel. AMPH induced significantly higher locomotion in *TH-Gal4/UAS-RicQ117L* flies than *TH-Gal4/+* flies at 5mM and 10mM AMPH doses (two-way ANOVA: Interaction: F_(3, 168)_ = 2.86, *p=0.04; Dose: F_(3, 168)_ = 13.48, ****p<0.0001; Genotype: F_(1, 168)_ = 22.22, ****p<0.0001; 5mM AMPH: **p=0.001, 10mM AMPH: **p=0.003, Sidak’s multiple comparisons test).

**Figure 4.**
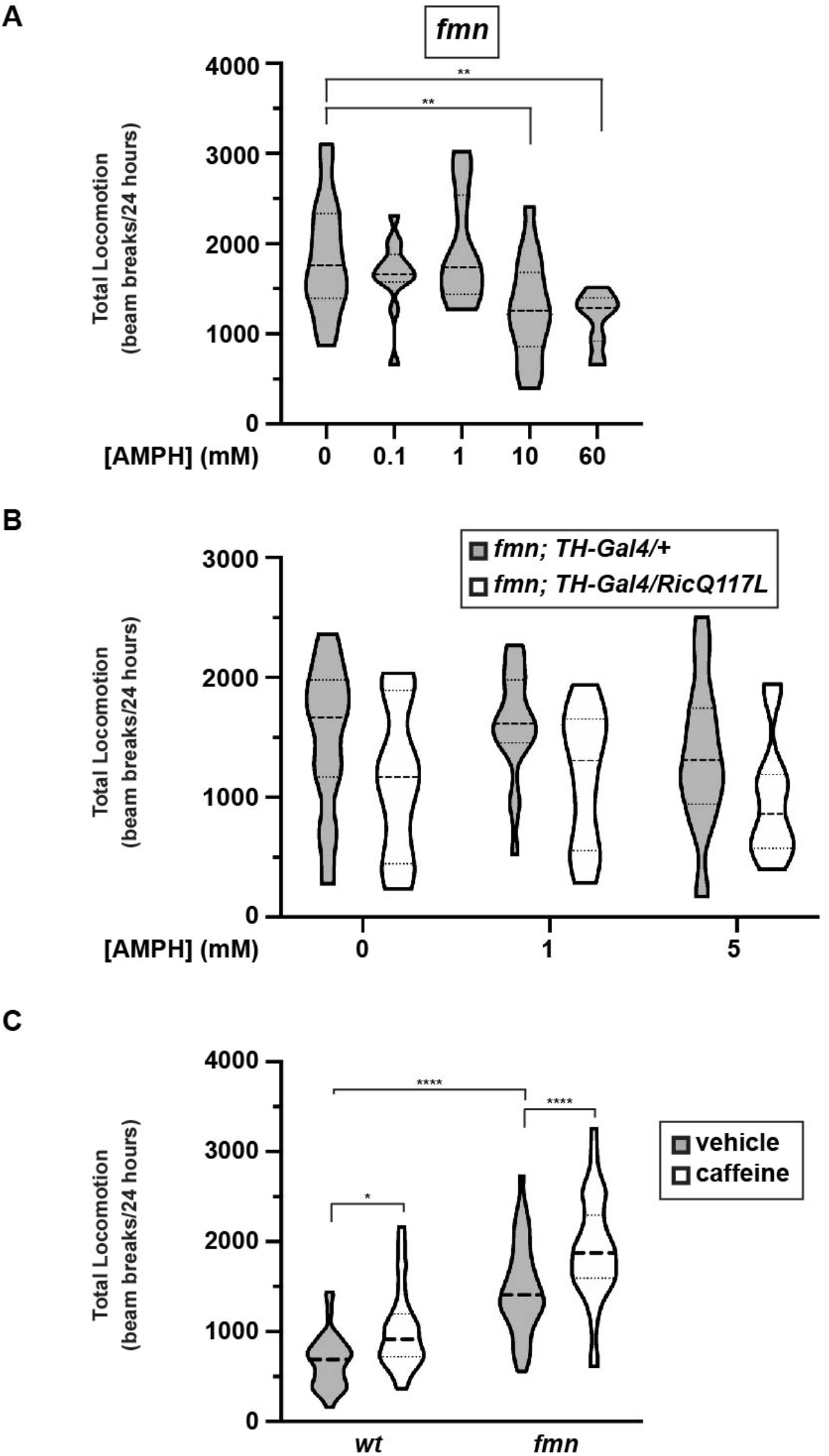
RicQ117L-enhanced AMPH sensitivity requires dDAT expression. *Locomotor response to AMPH feeding*. **(A)** *fmn fly response to AMPH. fmn* flies were fed either vehicle (n=21) or 0.1mM (n=13), 1.0mM (n=13), 10mM (n=16), or 60mM (n=15) AMPH, and locomotion was monitored as described in *Methods*. Data are presented as violin plots with median and quartiles indicated. AMPH had no significant on *fmn* locomotion, and significantly decreased locomotion at high doses as compared to vehicle-fed controls (one-way ANOVA F_(4, 70)_ = 5.895, p=0.0004; **p=0.007, ***p=0.004, Dunnett’s multiple comparisons test). **(B)** Male *fmn;TH-Gal4/+* (vehicle: n=24; 1.0mM: n=21; 5.0mM: n=23) and *fmn;UAS-RicQ117L* (vehicle: n=11; 1.0mM: n=9; 5.0mM: n=11) flies were fed either vehicle or the indicated AMPH doses and locomotion was measured as described in *Methods*. AMPH did not cause an interaction, or an effect of dose. There was a significant effect of genotype (two-way ANOVA: Interaction: F_(2, 95)_ = 0.057, p=0.95; Dose: F_(2, 95)_ = 1.43, p=0.25; Genotype: F_(1, 95)_ = 10.43, **p=0.002, n=10-24). There were no significant differences between the genotypes at either vehicle (p=0.24), 1mM (p=0.12), or 5mM (p=0.21) AMPH feeding conditions (Sidak’s multiple comparisons test). **(C)** *Locomotion in response to caffeine feeding. wt* (vehicle: n=42; caffeine: n=42) and *fmn* (vehicle: n=46; caffeine: n=42) fly locomotion was measured for 24 hours after feeding with either vehicle or 0.5 mg/ml caffeine, over 3 independent experiments. Data are presented as violin plots with median and quartiles indicated. Caffeine caused a significant effect of both drug and genotype (two-way ANOVA: Interaction: F_(1, 168)_ = 0.82, p=0.36; Drug: F_(1, 168)_ = 148.8, p<0.0001; Genotype: F_(1, 168)_ = 28.0, p<0.0001). *fmn* flies exhibited significantly more locomotion as compared to *wt* flies (p<0.0001), and caffeine significantly increased locomotion for both *wt* (p=0.013) and *fmn* (p<0.0001) flies (Tukey’s multiple comparisons test).

The enhanced AMPH sensitivity that RicQ117L imparts may be due to disrupted DAT trafficking, or due to an altered AMPH affinity for dDAT. To discern between these possibilities, we measured the ability of AMPH to inhibit DA uptake in HEK293T cells co-transfected with dDAT and either WT-Ric or RicQ117L. RicQ117L had no effect on the ability of AMPH to inhibit DA uptake (AMPH IC_50_: WT-Ric: 3.04±1.5µM vs RicQ117L: 4.1±2.3µM, p=0.70, two-tailed Student’s t test, n=6-7), suggesting that RicQ117L-enhanced AMPH sensitivity is likely due to Ric-dependent DAT trafficking, and not due to intrinsic changes in the AMPH/DAT interaction.

### Rit2 is required for AMPH-mediated DAT internalization in DAergic terminals

In addition to its effects on DA uptake and efflux, AMPH also stimulates robust DAT internalization in cell lines^67^, midbrain neurons^68,69^, and striatal DAergic terminals^24^. However, it is unknown whether either Rit2 or Ric are required for AMPH-stimulated DAT internalization, and it is possible that disrupted AMPH-mediated DAT trafficking may contribute to the observed increases in psychostimulant sensitivity following Rit2 and Ric manipulation in mice and flies, respectively. Given the lack of reagents to study native dDAT trafficking (there is no dDAT-specific antibody), we instead leveraged our previously reported approach to conditionally knock down Rit2 in mouse DAergic neurons^43,45^, and asked whether Rit2 is required for AMPH-stimulated DAT endocytosis in *ex vivo* dorsal striatum slices. As we previously reported, AMPH treatment significantly drove DAT internalization in dorsal striatal slices prepared from control-injected mice^24^. DAergic Rit2 silencing significantly attenuated AMPH-mediated DAT internalization, as compared to striata from mice injected with control AAV particles (Fig. 5). Thus, Rit2 is required for AMPH-stimulated DAT internalization, and disrupted AMPH-stimulated DAT trafficking may contribute to the enhanced psychostimulant sensitivity observed following Rit2 (and Ric) manipulation.

**Figure 5.**
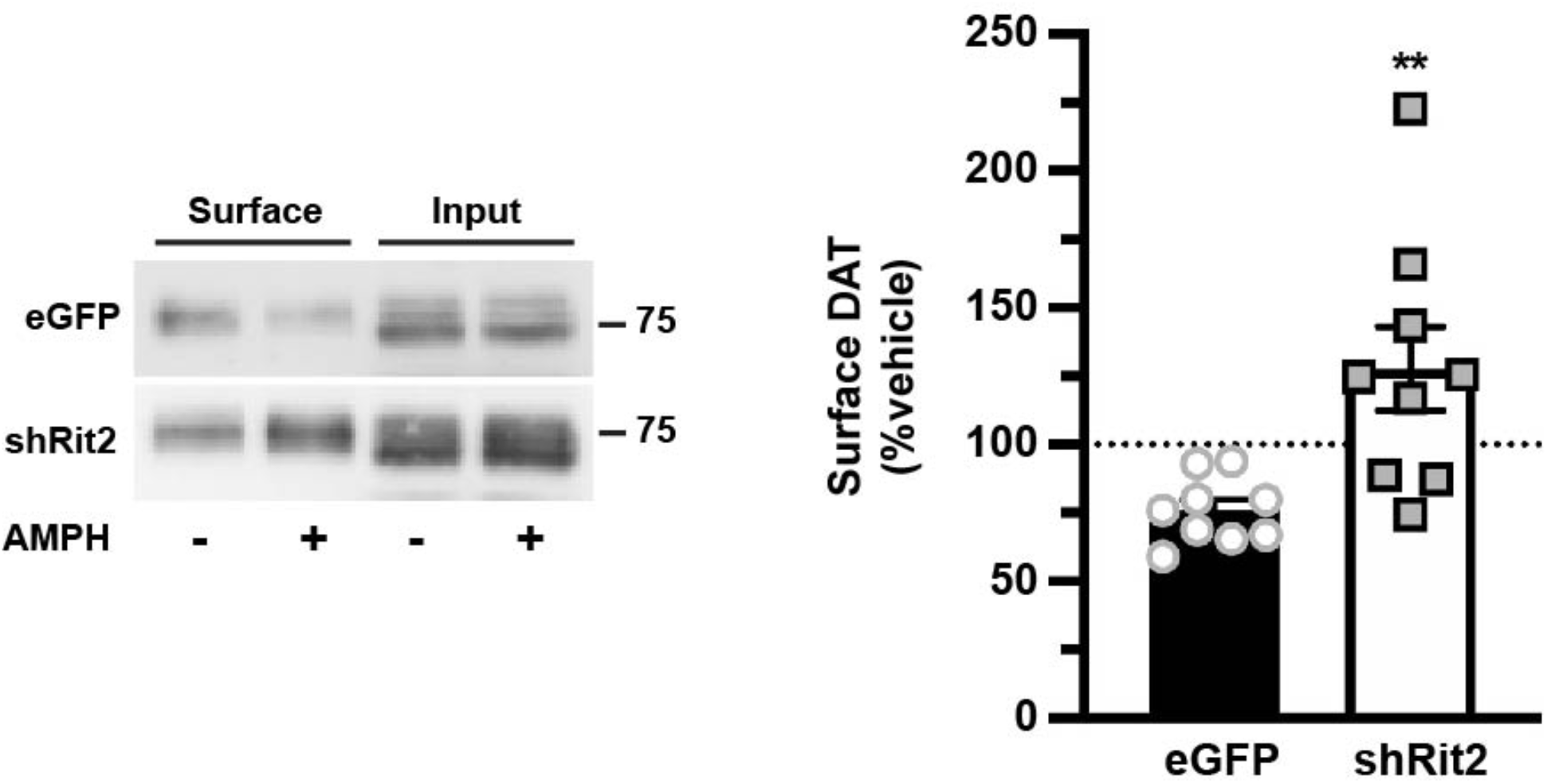
Rit2 is required for AMPH-stimulated DAT internalization in mouse striatum. *Ex vivo striatal slice biotinylation. Pitx3*^*IRES2-tTA*^ VTA were injected with AAV9 viral particles encoding either eGFP (n=7) or shRit2 (n=9), and acute striatal slices were prepared 4-6 weeks post-injection from 3 independent animals, as described in *Methods*. Contralateral hemi-slices were treated ±10µM AMPH, 30 min, 37°C, and surface DAT was isolated by biotinylation. *Left:* Representative DAT immunoblot. *Right:* Average surface DAT values, expressed as %vehicle-treated hemi-slice ±S.E.M. **p=0.005, one-tailed Student’s t test with Welch’s correction.

## DISCUSSION

The present study aimed to investigate how Rit2-dependent DAT trafficking impacts DA-dependent behaviors, using *Drosophila* as a model system. Given our current findings that dDAT and Ric interact, and that DAergic Ric activity impacts DAT activity, surface expression, baseline DA-dependent behaviors, and AMPH sensitivity, it is likely that *Drosophila* is an orthologous model for studying the DAT/Rit2 interaction. Indeed, *Drosophila* has proven a powerful system to investigate AMPH-stimulated DAT efflux mechanisms^15,47,48,65^, as well as hDAT coding variants identified in ADHD and ASD^21,49,70-72^. In the current study, *Drosophila* genetics enabled us to determine that Ric-mediated changes in behavior are, indeed, DAT-dependent, and not due to another Ric-mediated, DAT-independent mechanism. For instance, Ric activity was also demonstrated to stimulate neurite outgrowth in PC6 cells^73^. Ric binds to calmodulin^29^, and *in vivo* RicQ117L overexpression induced ectopic wing vein growth, which was exacerbated by concurrent null mutations in calmodulin^73^. Furthermore, genetic Ric ablation reduced fly viability in response to environmental stress^74^. Interestingly, a pan-neuronal RNAi screen identified Ric as a suppressor of olfactory-associated memory formation^75^, which requires DA signaling in the mushroom body^61,76^. Thus, our observations that DAergic Ric regulates other DA-dependent behaviors via its interaction with DAT are consistent with these reports.

What is the mechanism by which RicQ117L increases dDAT function and surface expression? We previously reported that DAT and Rit2 dissociate at the plasma membrane during PKC-stimulated DAT endocytosis^43,44^, and that the constitutively active Rit2 mutant lacked in its ability to dissociate from DAT following PKC activation^44^. Moreover, a SERT/DAT chimera, in which the DAT N-terminus was replaced with that of SERT, could not dissociate from Rit2, was resistant to PKC-stimulated endocytosis, and exhibited significantly decreased basal endocytosis^44^. Together these data raise the possibility that RicQ117L may inhibit dDAT/Ric dissociation, likewise stabilizing dDAT at the plasma membrane and increasing dDAT surface levels over time. However, it remains to be determined whether dDAT is subject to either PKC- or AMPH-stimulated endocytosis. Unfortunately, the excessive overexpression inherent in transient transfection is a significant obstacle to testing regulated dDAT trafficking *in vitro*, as we previously reported that regulated DAT trafficking is highly saturable and exquisitely sensitive to DAT expression level^77^. Moreover, the lack of dDAT-specific antibodies is prohibitive to studying dDAT trafficking *in situ*. Thus, additional reagents, and/or tagged fly lines, will be required to test whether dDAT undergoes regulated trafficking, and/or whether Ric activity plays a role in that process. However, the increased DAT surface expression *in vitro* and increased DAT activity *in situ* (Fig. 1) are consistent with a trafficking-mediated effect on DAT surface expression.

How might increased dDAT activity decrease sleep consolidation? Increases in DAT activity would be predicted to decrease extracellular DA half-life, leading to a shorter DA signal length. In addition, enhanced DA reuptake could increase the overall DAergic tissue content. Consistent with this possibility, extracellular DA concentrations are reduced by ∼50% in a DAT overexpression transgenic mouse, which expresses ∼2.5 times more total DAT protein^78^. This hypothesis is also consistent with the fact that *DAT*^*-/-*^ animals display 1) increased extracellular DA^9,79^, and 2) decreased sleep bout number (or increased wake episode length)^13,14^, as well as significantly reduced DAergic tone^9^. However, whether or not DA release and/or overall DA concentrations in the fly brain are altered by RicQ117L expression remain to be tested. dDAT is also required for sleep rebound characterized by increased sleep after mechanical sleep deprivation, indicating a role for dDAT in homeostatic sleep regulation^14^. Further, DAergic RicQ117L expression increased sleep bout frequency during the light phase of activity, but not at night (Fig. 2), suggesting a possible influence of circadian regulation. Extracellular DA concentrations and rates of DA uptake vary over the light/dark cycle, and DAT is required for circadian oscillations in DA release^80^. However, whether DAT is differentially sensitive to Ric-mediated regulation throughout the day remains unknown.

We previously reported that conditionally silencing Rit2 in mouse DA neurons decreased baseline anxiety, but had no effect on overall locomotion, consistent with our current results testing the role of DAergic Ric in *Drosophila* (Fig. S1 and S2). Notably, DAergic Rit2-KD altered acute cocaine responses, increasing cocaine sensitivity in male mice, and decreasing cocaine sensitivity in females^45^. Although cocaine binding to DAT is required for cocaine-induced reward^81^, it is unclear whether Rit2-KD alters the cocaine response though DAT. *Drosophila* also exhibit psychostimulant-mediated hyperactivity and preference^65,82,83^, and dDAT is required for AMPH-dependent hyperlocomotion^15^. Moreover, our current results indicate that DAergic CA Ric increases *Drosophila* sensitivity to AMPH DAT-dependently (Fig. 4), suggesting that DAT surface regulation impacts psychostimulant-induced hyperactivity. Interestingly, Rit2-KD and RicQ117L expression both increased psychostimulant sensitivity but had opposite effects on DAT surface expression. These data raise the possibility that disrupting Rit2/Ric GTPase by knockdown and mutant expression may lead to changes in DAT regulation that disrupt DA homeostasis, thereby increasing psychostimulant sensitivity, regardless of the absolute amount of surface DAT. Importantly, AMPH also stimulates DAT internalization in a Rho-dependent manner^69^. However, we do not know whether Rit2 acts with Rho to facilitate AMPH-induced DAT internalization, nor is it clear whether RIt2’s role in PKC- or AMPH-induced internalization are at play in the observed enhanced psychostimulant sensitivity. Nevertheless, future studies investigating the *in vivo* impact of regulated DAT trafficking, via leveraging DAT trafficking-dysregulated mutants, will more directly examine the impact of regulated DAT endocytosis on psychostimulant sensitivity and other DA-dependent behaviors.

## Acknowledgements

The authors wish to thank Tucker Conklin for excellent technical assistance, and Dr. Ratna Chaturvedi for assistance and training with *Drosophila* sleep studies and analyses.

