## Supplemental data for "Dopaminergic Ric GTPase activity impacts amphetamine sensitivity and sleep quality in a dopamine transporter-dependent manner in *Drosophila melanogaster*"

**Fagan et al**
**Supplemental data**

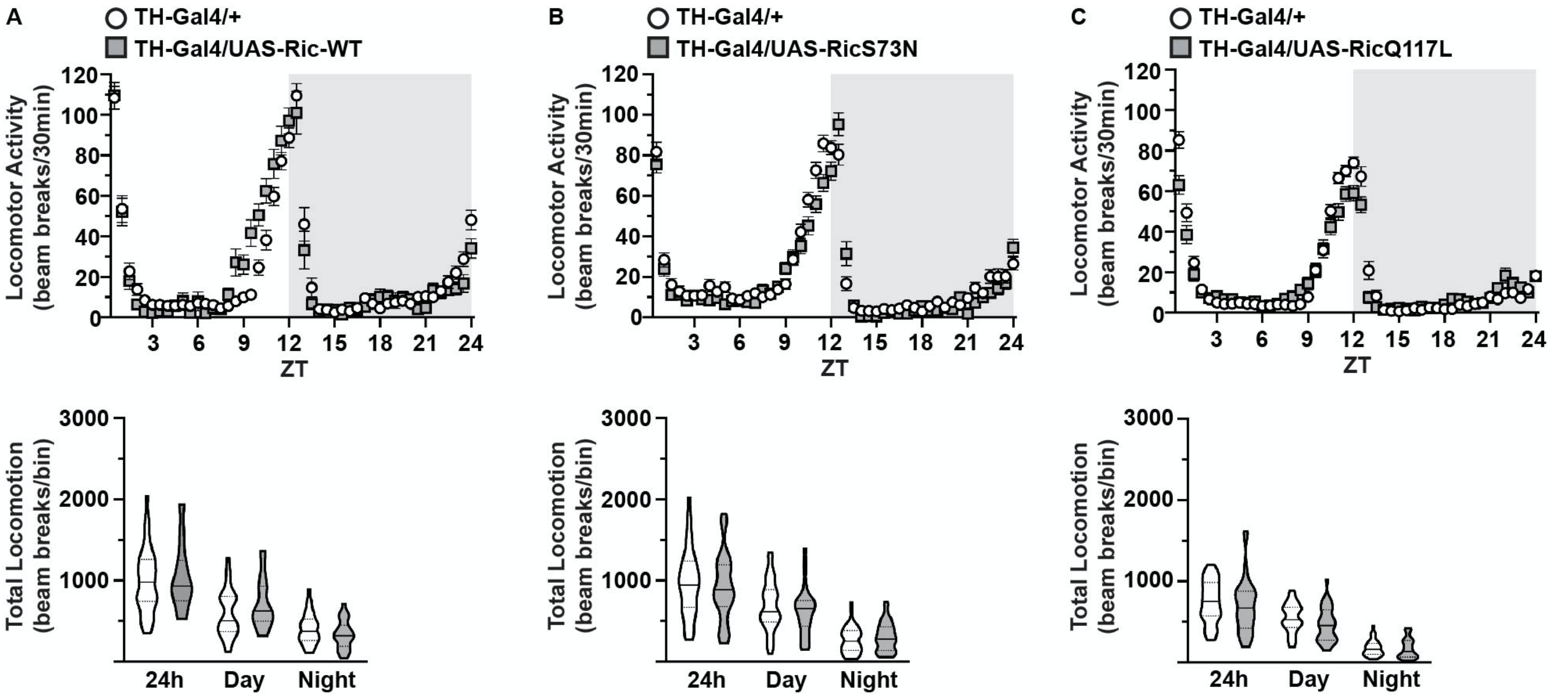


**Figure S1. Ric activity does not alter total *Drosophila* locomotor activity**. *TH-Gal4/+* (n=38), *TH-Gal4/UAS-Ric-WT* (n=24), *TH-Gal4/UAS-RicS73N* (n=26), and *TH-Gal4/UAS-RicQ117L* (n=36) fly locomotor activity was measured in progeny as the total number of beam breaks during 24h, day (12h lights-on), and night (12h lights-off) bins. Data are presented as violin plots indicating the median and quartiles.Data were analyzed by two-tailed student's *t* test or Welch's correction (as determined using Bartlett's test) between *TH-Gal4/+* control and experimental animals. Neither Ric-WT (**A**), RicS73N (**B**), nor RicQ117L (**C**) had an effect on total activity counts during the 24h, day, or night bins. Data were collected over 3-6 independent experiments.

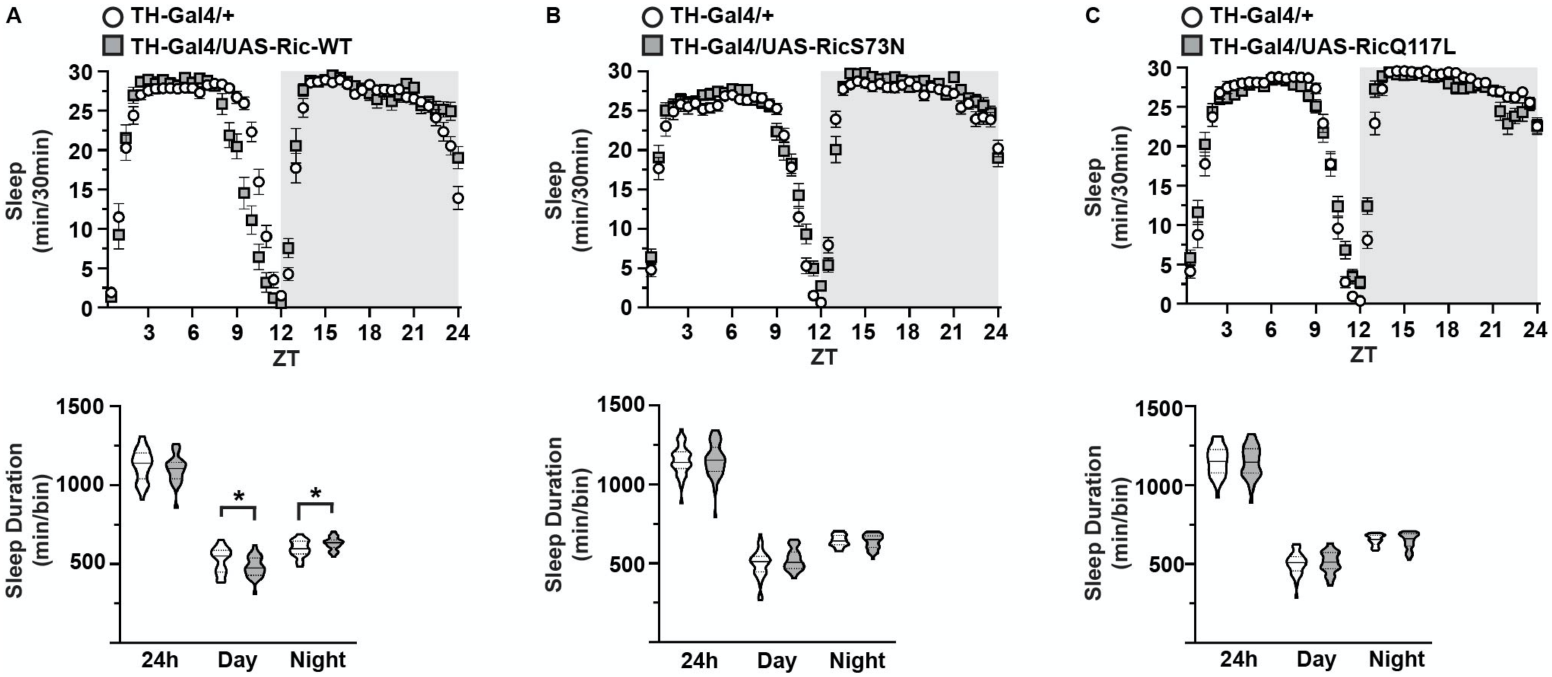


**Figure S2. Ric activity does not alter total *Drosophila* sleep.**
*TH-Gal4/+* (n=38), *TH-Gal4/UAS-Ric-WT* (n=24), *TH-Gal4/UAS-RicS73N* (n=26), and *TH-Gal4/UAS-RicQ117L* (n=36) fly total sleep was measured as the minutes spent sleeping during 24h, day (12h lights-on), and night (12h lights-off) bins. Data are presented as violin plots indicating the median and quartiles. Data were analyzed by two-tailed student's *t* test or Welch's correction (as determined using Bartlett's test) between *TH-Gal4/+* control and experimental animals. (**A**) Ric-WT had no effect on total sleep over the 24h period, but significantly decreased total sleep during the day (*p=0.03), and increased sleep during the night (*p=0.02). (**B**) RicS73N and (**C**) RicQ117L had no effect on total sleep in either the 24h, day, or night bins. Data were collected over 3-6 independent experiments.

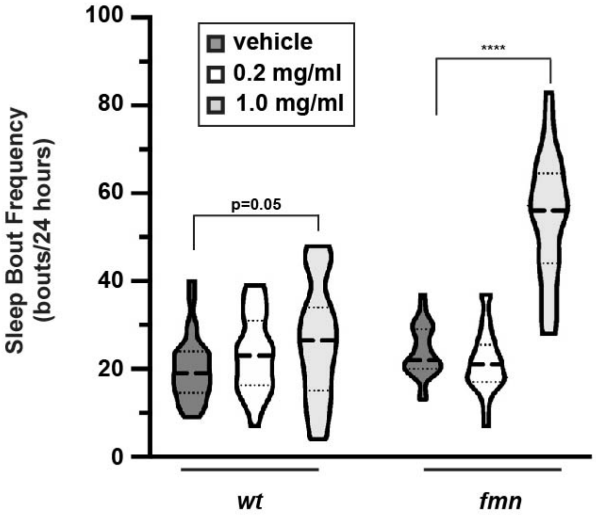


**Figure S3. Carbamazepine increases *fmn* sleep bout frequency.** *wt* (vehicle: n=29, CBZ-0.2mg/ml: n=28, CBZ-1.0mg/ml: n=28) and *fmn* (vehicle: n=28, CBZ-0.2mg/ml: n=26, CBZ-1.0mg/ml: n=30) fly sleep was measured over 24 hours during feeding with either vehicle or the indicated CBZ doses, beginning 24-hours following initial CBZ administration. Data are presented as violin plots indicating the median and quartiles. Asterisks indicate significant difference from vehicle control (two-way ANOVA: Interaction: $F_{(2, 158)} = 33.2$, $p<0.0001$; Dose: $F_{(2, 158)} = 57.94$, $p<0.0001$; Genotype: $F_{(1, 158)} = 41.0$, $p<0.0001$; n=26-30 flies assessed over 2 independent experiments). Sleep bout frequency was significantly increased by 1.0mg/ml CBZ for *fmn* ($p<0.0001$) and trended to increase for *wt* ($p=0.05$) as compared to vehicle-fed flies (Sidak's multiple comparison test).
